# System analysis of cross-talk between nuclear receptors reveals an opposite regulation of the cell cycle by LXR and FXR in human HepaRG liver cells

**DOI:** 10.1101/514976

**Authors:** Leonore Wigger, Cristina Casals-Casas, Michaël Baruchet, Khanh B. Trang, Sylvain Pradervand, Aurélien Naldi, Béatrice Desvergne

**Author notes:** Computational Systems Biology Team, Institut de Biologie de l’Ecole Normale Supérieure, Paris, France (Aurélien Naldi). The two first authors contributed equally (Leonore Wigger, Cristina Casals-Casas).

## Abstract

Transcriptional regulations exert a critical control of metabolic homeostasis. In particular, the nuclear receptors (NRs) are involved in regulating numerous pathways of the intermediate metabolism. The purpose of the present study was to explore in liver cells the interconnectedness between three of them, LXR, FXR, and PPARα, all three known to act on lipid and glucose metabolism, and also on inflammation. The human cell line HepaRG was selected for its best proximity to human primary hepatocytes. Global gene expression of differentiated HepaRG cells was assessed after 4 hours and 24 hours of exposure to GW3965 (LXR agonist), GW7647 (PPARα agonist), and GW4064 and CDCA (FXR synthetic and natural agonist, respectively). Our work revealed that, contrary to our expectations, NR specificity is largely present at the level of target genes, with a smaller than expected overlap of the set of genes targeted by the different NRs. It also highlighted the much broader activity of the synthetic FXR ligand compared to CDCA. More importantly, our results revealed that activation of FXR has a pro-proliferative effect and decreases polyploidy of hepatocytes, while LXR inhibits the cell cycle progression, inducing hepatocyte differentiation and a higher polyploidism. Conclusion: these results highlight the importance of analyzing the different NR activities in a context allowing a direct confrontation of each receptor outcome, and reveals the opposite role of FXR and LXR in hepatocyte cells division and maturation.

## Introduction

Homeostasis of energy metabolism results in a steady-state output of energy available for cell functions, despite the discontinuity of food intake and activities. Metabolic regulation in the liver is a major component of energy homeostasis. At the molecular level, metabolic regulation relies on three main types of control: allosteric, post-translational, and transcriptional. While most metabolic regulations benefit from a coordination of these mechanisms, transcriptional regulation exerts a critical control for keeping each component of the regulatory mechanisms at appropriate operating levels.

Nuclear receptors (NRs) are transcription factors that share many structural properties, notably a DNA binding domain folded in two zinc fingers and a ligand-binding pocket made of 13 alpha helices. Within the superfamily of NRs, which encompasses 48 members in humans, there is a sub-class called metabolic sensors. They are bound and activated by endogenous ligands that are metabolites belonging to the intermediary metabolisms, and actively contribute to the regulation of metabolic pathways. The discovery of each receptor initially emphasized the specificity of each receptor in a given metabolic pathway. For example, the peroxisome proliferator-activated receptors (PPARα, PPARβ/δ, PPARγ, also called NR1C1, NR1C2, NR1C3, according to the nomenclature agreed by the NC-IUPHAR Subcommittee on Nuclear Hormone Receptors) target genes in lipid metabolism, the farnesoid X receptors (FXR, also known as NRIH4) are involved in bile acid metabolism, and the liver X receptors (LXRα and LXRβ; NR1H3 and NR1H2, respectively) regulate cholesterol metabolism (1, 2). However, the classical linear view with each NR engaged in modulating one or a few pathways is challenged by the numerous and complex interconnections between the metabolism of glucide, lipids and amino acids, as well as by the numerous roles of NRs outside of metabolism. This highlights the need to delineate the regulatory network underlying homeostasis through systemic approaches.

The aim of this study was to explore the connections between the three NRs mentioned above. More specifically, PPARα is activated by unsaturated fatty acids and involved in many facets of both lipid and glucose metabolism. LXRα and LXRβ are activated by cholesterol derivatives but are also strongly lipogenic. Finally, FXR is bound by bile acids and is considered as a critical regulator of cholesterol metabolism (3). Thus, they clearly affect overlapping pathways. To better explore the interconnections, one must first assess the activity of each receptor in a given common and reproducible cellular context. For that purpose, we used the HepaRG hepatocarcinoma cell line, introduced in 2002 by Gripon et al. (4). HepaRG cells have been described as the closest in vitro model to human liver metabolism (5) and as a strong candidate for bioartificial liver applications (6).

Our work revealed that, in contrast to our expectations, NR specificity is largely present at the level of target genes, with a smaller than expected overlap of the set of genes targeted by the different NRs. The connection and coordination then mainly occur at the level of pathways. Our observations also highlight an important role of FXR and LXR in regulating cell growth and death of liver cells, with opposite effects on cell cycle progression and on hepatocyte polyploidization.

## Material and Methods

### Cell culture and chemicals

Cultures of HepaRG cells, generously provided by C. Guguen-Guillouzo, were grown as described in details in (4). After differentiation, hepatocyte-like cells were enriched by selective detachment using mild trypsinization and plated at 1.5×10^−5^/cm^2^ density in complete media plus 2% DMSO as described in (7). Based on pilot experiments, 2 μM for GW3965, 1 μM for GW4064 and GW7647 and 50 μM for CDCA were used. All compounds were diluted in DMSO, which was also used as vehicle control. The different agonists and CDCA were purchased from Sigma-Aldrich (St. Louis, MO).

### RNA extraction and Quantitative RT–PCR

Total RNA from cell culture was extracted with Trizol Reagent (Life Technologies, Carlsbad, CA), following the manufacturer’s instructions. The PCR arbitrary units of each gene were defined as the mRNA levels normalized to the *GAPDH* and the *EEF1A1* expression level in each sample using the qBase Software. The primers sequences are available upon request. The RNA for microarrays were prepared using: RNA quality assessment by Bioanalyzer chips; quantitative retrotranscription with iScript cDNA synthesis kit (Bio-Rad, Laboratories, Hercules, CA); real-time monitoring of PCR amplification of cDNA with FastStart Universal SYBR Green Master (Roche Applied Science, Indianapolis, IN) in an ABI Prism 7900 Sequence Detection System (Life Technologies, Carlsbad, CA).

### Microarrays

Microarrays were prepared from two time points, 4 hours and 24 hours, with 18 samples per time points: 3 samples of each of the 6 condition, i.e. FXR-L, LXR-L, PPARα-L, CDCA, vehicle controls (DMSO) and untreated controls. Samples were hybridized to Affymetrix GeneChip Human Gene 1.0 ST arrays (Affymetrix, Santa Clara, CA). Data summarisation and normalization was performed on all 36 samples together, using the robust multichip analysis (RMA) algorithm implemented in the Affymetrix Expression Console software. Probe sets that could not be mapped to any gene symbol were discarded. When more than one probe set was associated with the same gene symbol, only the probe set with the highest variance across the normalized expression levels in the 18 arrays from time point 4h was retained in the data set. After these two filtering steps, 20270 genes remained. Microarray data are available on GEO (accession number GSE124053).

### Differential Gene (DE) Expression

Differential gene expression for each treatment at each time point was analyzed with moderated t-tests performed with the bioconductor package LIMMA (8). Untreated and DMSO treated samples were combined in a single control group. For each time point, we applied a linear model with five experimental conditions (four treatments, one control) as coefficients. Four contrasts were extracted from the linear model, testing each of the four treatment conditions against the control condition. The resulting gene lists (limma output) are available in the supplementary data archive data_and_code.zip.

### Gene Set Enrichment Analysis (GSEA)

Gene set enrichment analysis (GSEA) was performed as described in (9) for each treatment at each time point. Most of the gene sets were taken from KEGG, while BIOCARTA sets from the Broad Institute’s Molecular Signatures Database (MsigDB version 3.0: www.broadinstitute.org/gsea/msigdb) are explicitly mentioned. Genes that were part of a gene set but that were not found on the Affymetrix chip via their gene symbol were removed from the gene sets. Of these reduced gene sets, those that had fewer than 10 or more than 500 genes were discarded, leaving a total of 327 gene sets. The gene sets were scored against the ranked lists from the LIMMA analysis (four per time point). The absolute t-values from the moderated t-tests were used for both ranking and weighting the genes. P-values were computed using sample permutation with 500 iterations, separately for each of the two time points. At each iteration, the limma model with four contrasts described in the section above was run on the permuted data in order to compute gene set scores for all four treatment-control comparisons. The estimated p-value for a gene set is the proportion of sample permutations that led to a higher score than the original data. To correct for multiple testing, the p-values of the gene sets were adjusted by the Benjamini-Hochberg method. This was done globally for the four gene set lists from time point 4 hours and for the four gene-set lists from time point 24h.

We considered a gene set potentially enriched if it had an adjusted p-value lower than 0.2. The leading edge subsets of significantly enriched pathways were extracted, using the GSEA algorithm. The resulting lists of pathways are available in the supplementary data archive supplementary_archive.zip.

### Cell-cycle analysis

As described in (10), cells were fixed in 70% ice-cold ethanol, washed in PBS and resuspended in staining solution containing 20 μg/mlpropidiumiodide,0.05%TritonX-100g/ml propidium iodide, 0.05% Triton X-100 and 200 μg/ml propidium iodide,0.05%TritonX-100g/ml RNAse A (Sigma-Aldrich, St. Louis, MO) and 10,000 cells were analyzed per sample using the BD bioscience FACSDiva. Data were based on 8 replicates from 3 independent experiments.

### Western Blot

SDS-PAGE was performed using cultured HepaRG cell extracts and transferred to nitrocellulose membranes (Novex, Life Technologies, Carlsbad, CA). After blocking, bands were detected with primary antibodies to p27 (#2552), CCND3 (#2936) and RB (RB Antibody Sampler Kit #9969) and secondary antibodies, anti-rabbit (#7074) or anti-mouse (#7076) IgG conjugated to horseradish peroxidase (Cell Signaling Technology Inc., Danvers, MA), and applying the SuperSignal^®^ West Pico Chemiluminescent Substrate (Thermo Fisher Scientific, Bremen, Germany). Membranes were then exposed to x-ray films (GE Healthcare, Milwaukee, WI). Beta-actin(#A2228,Sigma-Aldrich,St.Louis,MO)wasusedasaloadingeta-actin (#A2228, Sigma-Aldrich, St. Louis, MO) was used as a loading control.

## Results

### Defining and validating the experimental set-up

The aim of the experiment was to evaluate which hypothesis could best explain the cross-regulatory action of the nuclear receptors on metabolic pathways. Do the receptors regulate common target genes, or do they regulate different sets of genes belonging to common pathways?

Differentiated HepaRG cells were chosen since their global gene expression patterns are much closer to liver or primary human hepatocyte cultures than the widely used HepG2 cell line (5). The mRNA expression levels of PPARα, LXRα/β and FXR in HepaRG cells are highly consistent with those observed in human primary liver cells. To activate each receptor, we used synthetic ligands known to be the most specific: GW4064 for FXR, GW3965 for LXRs, and GW7647 for PPARα, hereafter called FXR-L, LXR-L, and PPARα-L. We also used the bile acid CDCA, which is a natural ligand of FXR (11, 12). CDCA is also known to affect LXR (13), PPARα (14) and HNF4 (15) pathways, even though the mechanisms are not yet well understood. For each compound, we selected the lowest concentration that triggered a good response of canonical targets in a pilot experiment (data not shown).

Microarrays were run on differentiated HepaRG first prior any treatment, then 4 hours after the beginning of the cell exposure to each treatment to identify direct target genes, and finally 24h after the beginning of the treatment to identify global crosstalk. We first verified that the NRs of interest were indeed well expressed in HepaRG cells. Next, we carried out a differential expression analysis, comparing ligand-treated versus control samples (CDCA, FXR-L, LXR-L or PPARα compared to controls) at each of the two time points (4h and 24h). We identified the genes that were significantly affected by any ligand treatment at either time point and used their mRNA expression data as features to perform a hierarchical clustering of the five different cellular models analyzed by Hart *et al*. (5), as explained in supplemental Figure 1. The profile of our set of DE genes was found to be similar in human liver and in primary hepatocytes, whereas HepG2 cells have a gene expression pattern far from all the other models analyzed. The expression profile of HepaRG cells (differentiated and undifferentiated) was distinct from that of liver cells, but much closer to it than that of HepG2 cells.

Altogether these initial observations confirmed that HepaRG cells are, in this context, more pertinent than other *in vitro* models widely used in literature, such as HepG2 cells (16, 17).

### NR cross-talks happen at the pathway level and are mostly related to metabolism

The analyses of the set of genes differentially expressed at the 4 hours time point identified 681 genes affected by at least one treatment. A good part of these genes are associated to the FXR-L treatment (~500 genes) and to a lesser extent to CDCA (~160 genes), while fewer are associated to PPARα-L (~90), and to LXR-L (<10) (Fig 1A, left part). This important variation in the number of genes modulated by each treatment is consistent with that shown in studies analyzing gene expression or ChIP-seq specifically for one or the other NR (18–20). These differentially expressed genes include the main known targets of the selected nuclear receptors. For example, FXR-L and CDCA modulate *ABCB11* (*BSEP*), *ABCB4* (*MDR3*), *CYP7A1* and *PGC1* expression. Some known FXR target genes are found modulated only by FXR-L, and not by CDCA, such as *NR0B2* (*SHP*), *HNF4A* and *PPARα*. Others are only regulated by CDCA, including *UGTB4, RORA, HMGCS1* and *NR1D1*/2. PPARα-L up-regulates its well-established target genes *FABP4*, *HMGCS2, ACOX1, CD36* and *CPT1A*. LXR-L has a very modest effect, modulating less than 10 genes, which include the known LXR target genes *SCD, SREBF1* (also called *SREBP*), *ABCA1, FASN*, and *MYLIP/IDOL*.

**Figure 1:**
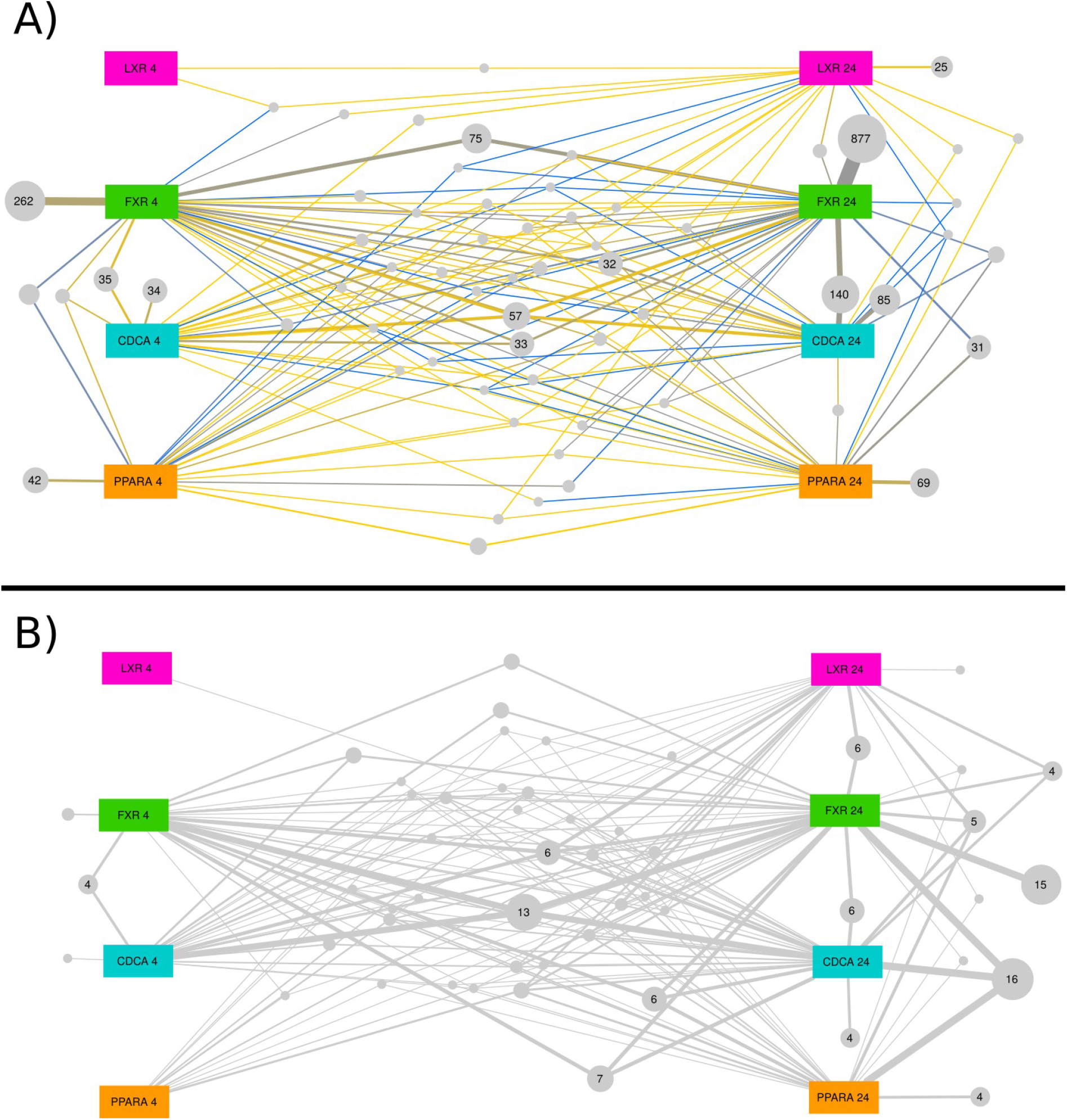
Visual Representation of the specific and shared effects of the treatments. Large colored nodes denote the 4 treatments at 4h (on the left) or at 24h (on the right), while grey bubbles denote groups of regulated genes (panel A) or of enriched pathways (panel B), which are associated to the same set of treatments. For example, panel A shows that 1214 genes are regulated only by FXR-L: 262 only after 4h, 75 at both time-points and 877 only at 24h. Similarly, the center bubble indicates that 57 genes are regulated by both FXR-L and CDCA at both time-points. The size of the bubbles and of the connections with their associated treatments is proportional to the number of genes or pathways in the group. In panel A, the color of the connection further indicates the ratio between genes which are activated (yellow) or inhibited (blue) by this treatment. The original Cytoscape files are available as supplementary material and a summary of the observed overlap is available in Table 1.

We then identified 1611 differentially expressed genes at the 24h time-point (Fig 1A, right part). This strong increase in this number of genes, compared to the early time-point, is consistent with the identification of indirect effects. For example, *SHP* (*NR0B2*), a well-described FXR target gene very significantly increased at 4 hours by FXR-L, mediates indirect regulations, such as the strong down-regulation of CYP7A1 seen at 24h (21). Also, SREBF, which is induced by LXR at 4h, is a regulator of key genes in lipid metabolism (22). These genes are in turn induced at 24h, as for example *SCD1*, *ACLS1*, *HMGCR* and *HMGCS1*. Like at 4h, most genes were associated to FXR-L (~1100) and CDCA (~350) and fewer to PPARα-L (~140) and LXR-L (<50).

Intriguingly, less than half of the genes identified at 4h were still differentially expressed at 24h (312 out of 681) (Table 1), suggesting that most direct effects are only transient. Comparing the different treatments shows surprisingly few cross-talks at the level of transcriptional regulation: among the 1980 differentially expressed genes, 75% are specific to a single treatment. As expected, FXR-L and CDCA have the biggest overlap, sharing 63% of the common genes (16% of all the differentially expressed ones). Nevertheless some genes are common targets for different NRs. For example, *PDK4, HMGCS2*, and *ACOX1*, involved in lipid metabolism, are all up-regulated by PPARα-L and down-regulated by FXR-L; *ABCA1* and *MYLIP*, well described target genes of LXR (3, 23), are also down-regulated by FXR. Some other genes involved in lipid and glucose metabolism, such as *FASN, FABP4, LPL, INSIG1, SCD*, are found up-regulated both by PPARα-L and LXR-L (Fig. 1A and Table 1). To explore the possibility of cross-talk at the level of pathways, we looked for enriched pathways shared across time-points and treatments. We identified 153 enriched pathways in our dataset. Like for differentially expressed genes, we observed an increase in the number of enriched pathways between the early and late time-points. Most enriched pathways (54%) were identified at both time-points (compared to 16% of the differentially expressed genes). Even though they remain predominantly associated to the FXR-L (140) or CDCA (114) treatments, PPARα-L (62) and LXR-L (46) treatments are also well represented. Importantly, 80% (123/156) of the enriched pathways are shared between several treatments, compared to 25% (500/1980) at the gene level. While 40 of the 123 shared pathways are still specific to the related FXR-L and CDCA treatments, which both target the same NR, the remaining 83 pathways are associated with two or even three treatments targeting separate NRs and thus point to cross-talk at the pathway level. In other words, more than half of all enriched pathways (83/153) are associated with at least two NRs (Figure 1B; Table 1).

**Table 1:** Summary of the distribution of differentially expressed (DE) genes and enriched pathways among time points and treatments. Top four rows: number and percentage of DE genes and positively enriched KEGG pathways from all treatments combined, found either at a single time point (4h, 24h) or at both time points (4h and 24h); totals from both time points. Remaining table: Numbers and percentages found per ligand treatment (at any or both time points), with a breakdown of how many DE genes and enriched pathways are exclusive to one treatment and how many are shared between two or multiple treatments. This summary shows that cross-talks observed at the pathway level are not visible when looking solely at differentially expressed genes.

|  |  | Genes |  | KEGG Pathways |  |
| --- | --- | --- | --- | --- | --- |
|  |  | # | % | # | % |
| All ligands combined | 4h | 407 | 20.35 | 7 | 4.58 |
|  | 24h | 1264 | 63.20 | 64 | 41.83 |
|  | 4h and 24h | 329 | 16.45 | 82 | 53.59 |
|  | Total | 2000 |  | 153 | 100.0 |
| CDCA-L | alone | 119 | 23.02 | 5 | 4.39 |
|  | shared with FXR-L | 318 | 61.51 | 40 | 35.09 |
|  | shared with LXR-L | 7 | 1.35 | 1 | 0.88 |
| | shared with PPARα-L | 6 | 1.16 | 0 | 0.00 |
|  | shared with multiple | 67 | 12.96 | 68 | 59.65 |
|  | Total | 517 | 100.00 | 114 | 100.00 |
| FXR-L | alone | 1214 | 71.08 | 20 | 14.29 |
|  | shared with CDCA | 318 | 18.62 | 40 | 28.57 |
|  | shared with LXR-L | 14 | 0.82 | 7 | 5.00 |
| | shared with PPARα-L | 91 | 5.33 | 4 | 2.86 |
|  | shared with multiple | 71 | 75.23 | 69 | 63.57 |
|  | Total | 1708 | 100.00 | 140 | 100.00 |
| LXR-L | alone | 26 | 38.81 | 1 | 2.17 |
|  | shared with CDCA | 7 | 10.45 | 1 | 2.17 |
|  | shared with FXR-L | 14 | 20.90 | 7 | 15.22 |
| | shared with PPARα-L | 6 | 8.96 | 1 | 2.17 |
|  | shared with multiple | 14 | 59.70 | 36 | 80.43 |
|  | Total | 67 | 100.00 | 46 | 100.00 |
| PPARα-L | alone | 127 | 43.05 | 4 | 6.45 |
|  | shared with CDCA | 6 | 2.03 | 0 | 0.00 |
|  | shared with FXR-L | 91 | 30.85 | 4 | 6.45 |
|  | shared with LXR-L | 6 | 2.03 | 1 | 1.61 |
|  | shared with multiple | 65 | 22.03 | 53 | 85.48 |
|  | Total | 295 | 100.00 | 62 | 100.00 |

The most striking observation at this point is that few genes are co-regulated by the NRs studied herein. In contrast, the cross-talk appeared in the associated pathways. We therefore explored further the shared enriched pathways to discover new NR cross-talks.

### Cell growth and death is overrepresented in FXR and LXR cross-talks

Based on the KEGG pathway database (www.genome.jp/kegg/pathway.html), we manually classified enriched pathways into 4 groups: “Metabolism”, “Cell Growth and Death”, “Immune System” and “Other”. ‘Transport and catabolism’, ‘Endocrine, Digestive, Excretory Systems’ and ‘Endocrine and Metabolic diseases’ were merged into the ‘Metabolism’ group. ‘Genetic Information Processing’, ‘Cellular Processes’ (other than ‘Peroxisome’) and ‘Cancers’ form the ‘Cellular Growth and Death’ group. Finally, ‘Immune and Infectious Diseases’ correspond to our ‘Immune System’ group. Unclassified pathways that we could not associate to one of these 3 groups form the ‘Other’ group (the names of the pathways in each category are made available in the supplementary archive “data_and_code.zip”).

As a set of core genes may drive different pathways belonging to the same group, we then counted the genes belonging to the leading edge of the GSEA curves of several pathways within the same group (illustrated in supplementary Figure S2). In the metabolism group, only 14 genes are involved in at least 10% of the pathways. This amount increases for the two other groups with 121 genes being involved in at least 10% of pathways of cell growth and death and 96 for the immune system; 37 of them are common to both groups. As most DE genes are specific to one pathway, we are confident about the relevance of the enriched pathways.

The ‘Metabolism’ group covers 48% of the enriched pathways, ‘Cellular Growth and Death’ covers 30%, and ‘Immune System’ 14%. Unsurprisingly, all pathways shared by the three NR and 65% of the ones shared by FXR and PPARα are involved in metabolism (Table 2). This analysis confirmed that the largest group of enriched pathways with the highest specificity of involved genes concerns metabolic regulations.

**Table 2:** Distribution of categories of enriched pathways depending on their association with treatments. Enriched KEGG pathways were grouped manually into 4 wide categories (“Metabolism”, “Cell growth and death”, “Immune system” and “Other”). Top four rows: Number and percentage of positively enriched KEGG pathways from all treatments combined, found either at a single time point (4h, 24h) or at both time points (4h and 24h) in each category; totals from both time points. Remaining table: Numbers and percentages of pathways found per ligand treatment (at any or both time points), with a breakdown of how many DE genes and enriched pathways are exclusive to one treatment and how many are shared between two or multiple treatments. Bottom section: Numbers and percentages of pathways found if the pathways lists from CDCA and FXR-L treatments are merged.

|  |  | KEGG Pathways: Metabolism |  | KEGG Pathways: Cell Growth |  | KEGG Pathways: Immune |  | KEGG Pathways: Others |  |
| --- | --- | --- | --- | --- | --- | --- | --- | --- | --- |
|  |  | # | % | # | % | # | % | # | % |
| All ligands combined | 4h | 3 | 4.17 | 1 | 2.08 | 3 | 13.04 | 0 | 0.00 |
|  | 24h | 38 | 52.78 | 17 | 35.42 | 7 | 30.43 | 2 | 20.00 |
|  | 4h and 24h | 31 | 43.06 | 30 | 62.50 | 13 | 56.52 | 8 | 80.00 |
|  | Total | 72 | 100.00 | 48 | 100.00 | 23 | 100.00 | 10 | 100.00 |
| CDCA-L | alone | 1 | 1.85 | 3 | 8.11 | 1 | 6.67 | 0 | 0.00 |
|  | shared with FXR-L | 9 | 16.67 | 14 | 37.84 | 10 | 66.67 | 7 | 87.50 |
|  | shared with LXR-L | 1 | 1.85 | 0 | 0.00 | 0 | 0.00 | 0 | 0.00 |
| | shared with PPARα-L | 0 | 0.00 | 0 | 0.00 | 0 | 0.00 | 0 | 0.00 |
|  | shared with multiple | 43 | 79.63 | 20 | 54.05 | 4 | 26.67 | 1 | 12.50 |
|  | Total | 54 | 100.00 | 37 | 100.00 | 15 | 100.00 | 8 | 100.00 |
| FXR-L | alone | 6 | 9.23 | 7 | 15.91 | 6 | 28.57 | 1 | 10.00 |
|  | shared with CDCA | 9 | 13.85 | 14 | 31.82 | 10 | 47.62 | 7 | 70.00 |
|  | shared with LXR-L | 3 | 4.62 | 3 | 6.82 | 1 | 4.76 | 0 | 0.00 |
| | shared with PPARα-L | 3 | 4.62 | 0 | 0.00 | 0 | 0.00 | 1 | 10.00 |
|  | shared with multiple | 44 | 67.69 | 20 | 45.45 | 4 | 19.05 | 1 | 10.00 |
|  | Total | 65 | 100.00 | 44 | 100.00 | 21 | 100.00 | 10 | 100.00 |
| LXR-L | alone | 1 | 3.45 | 0 | 0.00 | 0 | 0.00 | 0 | N/A |
|  | shared with CDCA | 1 | 3.45 | 0 | 0.00 | 0 | 0.00 | 0 | N/A |
|  | shared with FXR-L | 3 | 10.34 | 3 | 20.00 | 1 | 50.00 | 0 | N/A |
| | shared with PPARα-L | 1 | 3.45 | 0 | 0.00 | 0 | 0.00 | 0 | N/A |
|  | shared with multiple | 23 | 79.31 | 12 | 80.00 | 1 | 50.00 | 0 | N/A |
|  | Total | 29 | 100.00 | 15 | 100.00 | 2 | 100.00 | 0 | N/A |
| PPARα-L | alone | 2 | 4.26 | 1 | 11.11 | 1 | 25.00 | 0 | 0.00 |
|  | shared with CDCA | 0 | 0.00 | 0 | 0.00 | 0 | 0.00 | 1 | 50.00 |
|  | shared with FXR-L | 3 | 6.38 | 0 | 0.00 | 0 | 0.00 | 0 | 0.00 |
|  | shared with LXR-L | 1 | 2.13 | 0 | 0.00 | 0 | 0.00 | 0 | 0.00 |
|  | shared with multiple | 41 | 87.23 | 8 | 88.89 | 3 | 75.00 | 1 | 50.00 |
|  | Total | 47 | 100.00 | 9 | 100.00 | 4 | 100.00 | 2 | 100.00 |
| CDCA, FXR-L (merged) | alone | 16 | 32.65 | 24 | 51.06 | 17 | 77.27 | 8 | 80.00 |
|  | shared with LXR-L | 8 | 16.33 | 15 | 31.91 | 2 | 9.09 | 0 | 0.00 |
| | shared with PPARα-L | 25 | 51.02 | 8 | 17.02 | 3 | 13.64 | 2 | 20.00 |
|  | Total | 49 | 100.00 | 47 | 100.00 | 22 | 100.00 | 10 | 100.00 |

However, 72% of the pathways associated to both FXR-L and LXR-L are related to cell growth and death, while this pathway group does not represent more that 30% in the other combinations of treatments. In the literature, the role of FXR and LXR in liver is considered as mostly related to metabolism regulation with main roles in lipid, cholesterol and glucose metabolism, FXR being anti-lipogenic and LXR pro-lipogenic (1). However, they also play an important role in liver regeneration (24–26) or cancer development (27, 28), which fits with our finding that their cross-talk at pathway level is to a large extent related to cell growth and death.

### FXR and LXR oppositely regulate the cell cycle progression

To understand the mechanisms involving FXR and LXR in the regulation of cell growth and death, we focused on the “Cell cycle” pathway, which is enriched in four conditions: at 4h by FXR-L and CDCA, and at 24h by FXR-L and LXR-L. We further looked at the genes that belong to the leading edges of the cell cycle pathway at 24h for FXR-L (45 genes) and LXR-L (41 genes). Intriguingly, FXR-L predominantly activates gene expression while LXR-L inhibits it. Twenty-three of the 26 genes found in the leading edges for both treatments are oppositely regulated by FXR-L and LXR-L (Fig. 2).

**Figure 2:**
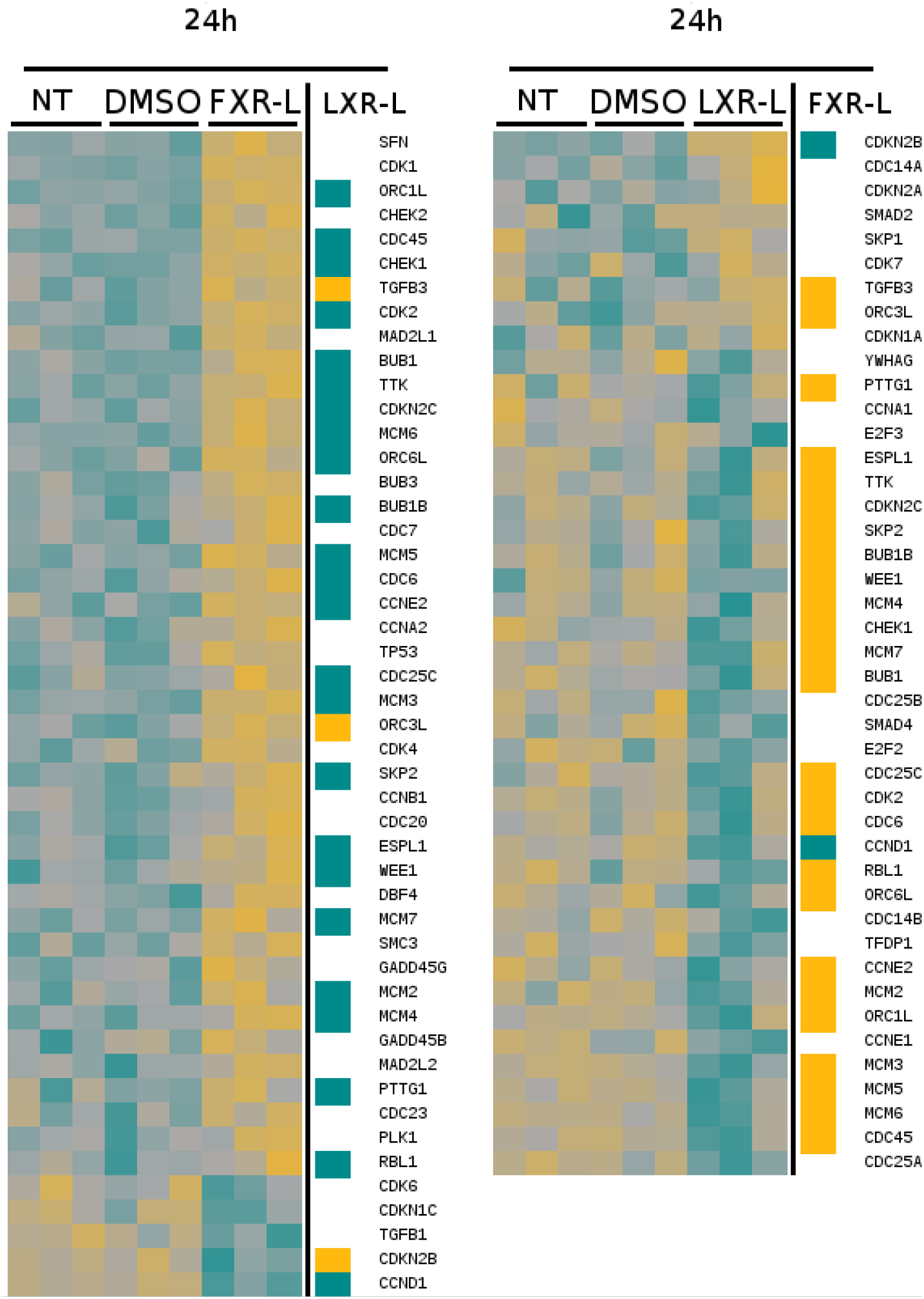
Effect of FXR-L and LXR-L treatments on genes associated to the cellcycle pathway. These heatmaps show the expression levels of genes driving the cell-cycle pathway at 24h for the FXR-L (on the left) and LXR-L (on the right) treatments compared to the controls (non-treated (NT) and DMSO-treated) according to GSEA (see methods and supplementary material). For each gene, the 9 experimental values are centered around their mean (in grey). Larger values appear in yellow, while lower values are blue. Each heatmap has an extra column on the right showing the tendency of the common genes for the other treatment. With FXR-L treatment, most genes are up-regulated, while they are down-regulated with LXR-L. Among the 27 genes regulated by the two treatments, only 3 have the same profile in the two conditions (TGFB3, ORC3L and CCND1).

Using the KEGG diagrams to illustrate the corresponding pathway and genes (Fig. 3 and supplementary Fig. S3), the patterns of FXR-L at 4h show an up-regulation of genes involved in the G1/S transition like *C-MYC*, and *CCND1*. At 24h, the cell cycle key players are upregulated by FXR-L and downregulated by LXR-L. The list comprises *CCNE, CDK2*, and *P18* as well as genes involved in DNA replication such as *CDC6* and members of the minichromosome complex (*MCM*) (29). These observations suggest a pro-proliferative role for FXR, with an up-regulation of genes involved in the G1/S progression at 4h and a more global positive regulation at 24h. LXR have the opposite effect inhibiting genes involved in the cell cycle progression and DNA replication.

**Figure 3:**
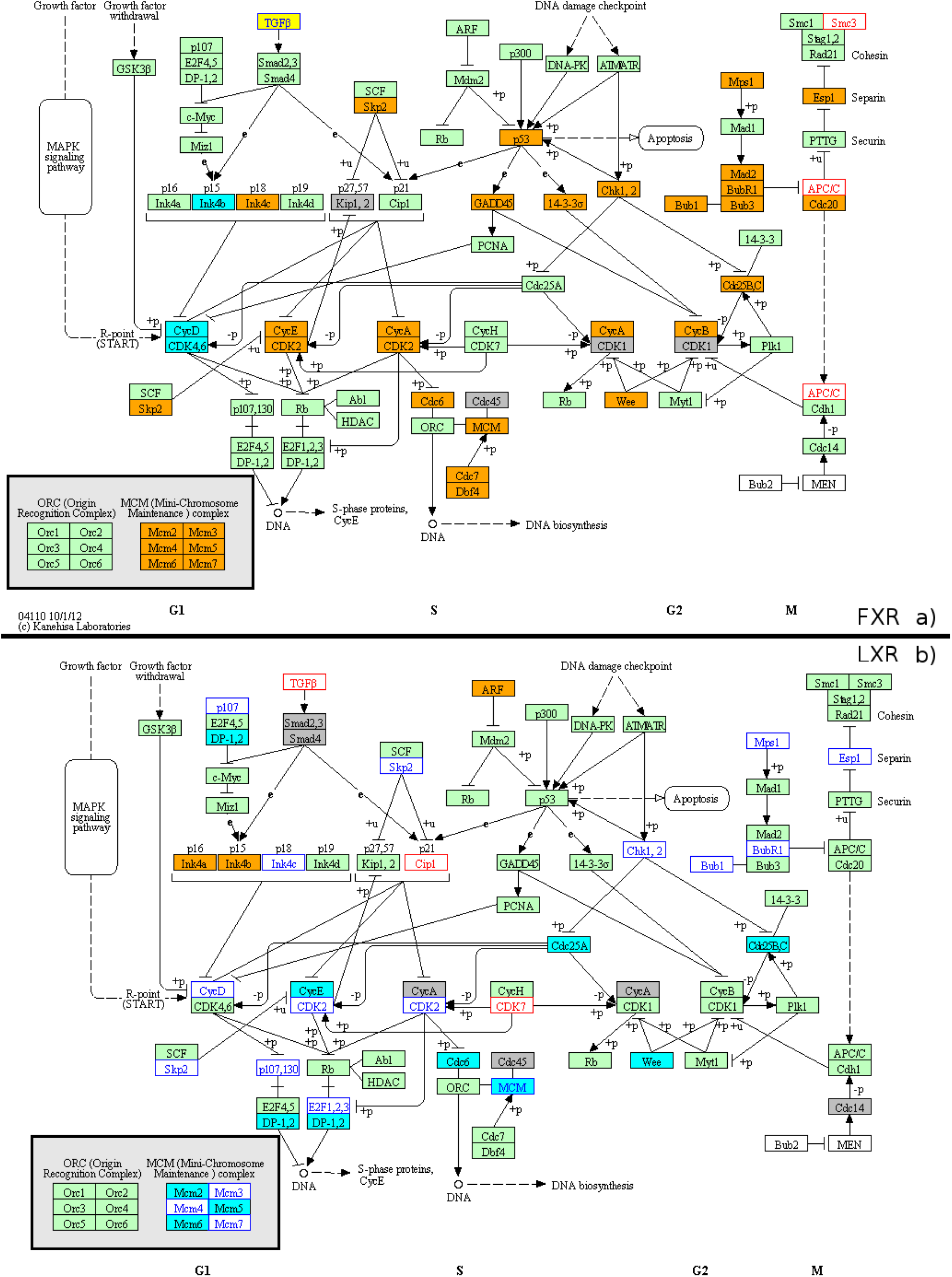
Visualisation of the effects of FXR-L and LXR-L on the KEGG cell cycle pathway map. Genes found in the leading edge of the cell cycle pathway at 24h after FXR-L (panel A) and LXR-L (panel B) treatments are highlighted. Boxes with a blue (resp. orange) background denote down-regulated (resp. up-regulated) genes (|log foldchange| > 1.2). White boxes with blue (resp. red) text denote smaller down-regulations (resp. upregulations). The yellow TGFβ box in panel A corresponds to the TGFβ1, TGFβ2 and TGFβ3 genes which are regulated in opposite directions.

To determine if these gene modulations reflect an actual regulation of the cell cycle, we analyzed the distribution of cell cycle phases by propidium iodide staining and flow cytometry. We treated HepaRG cells with FXR-L and LXR-L for 24, 48 or 72 hours. Knowing the pro-proliferative effect of serum in the culture media, the experiment was made both in the presence and in the absence of serum. Flow cytometry analyses revealed two well separated populations corresponding to diploid and tetraploid populations (2C and 4C DNA content, respectively), characteristic of mammalian hepatocytes (Fig. 4A) (30). In serum free conditions, FXR-L treatment triggered a decrease of the tetraploid cell population, visible at 24h and significant at 48h and 72h, with 25% and 36% less 4C cells respectively, while LXR-L treatment has no significant effect (Fig. 4D). In the presence of serum, the 4C population also decreases at 72h with FXR-L (22%), whereas it significantly increases with LXR-L (17%).

**Figure 4:**
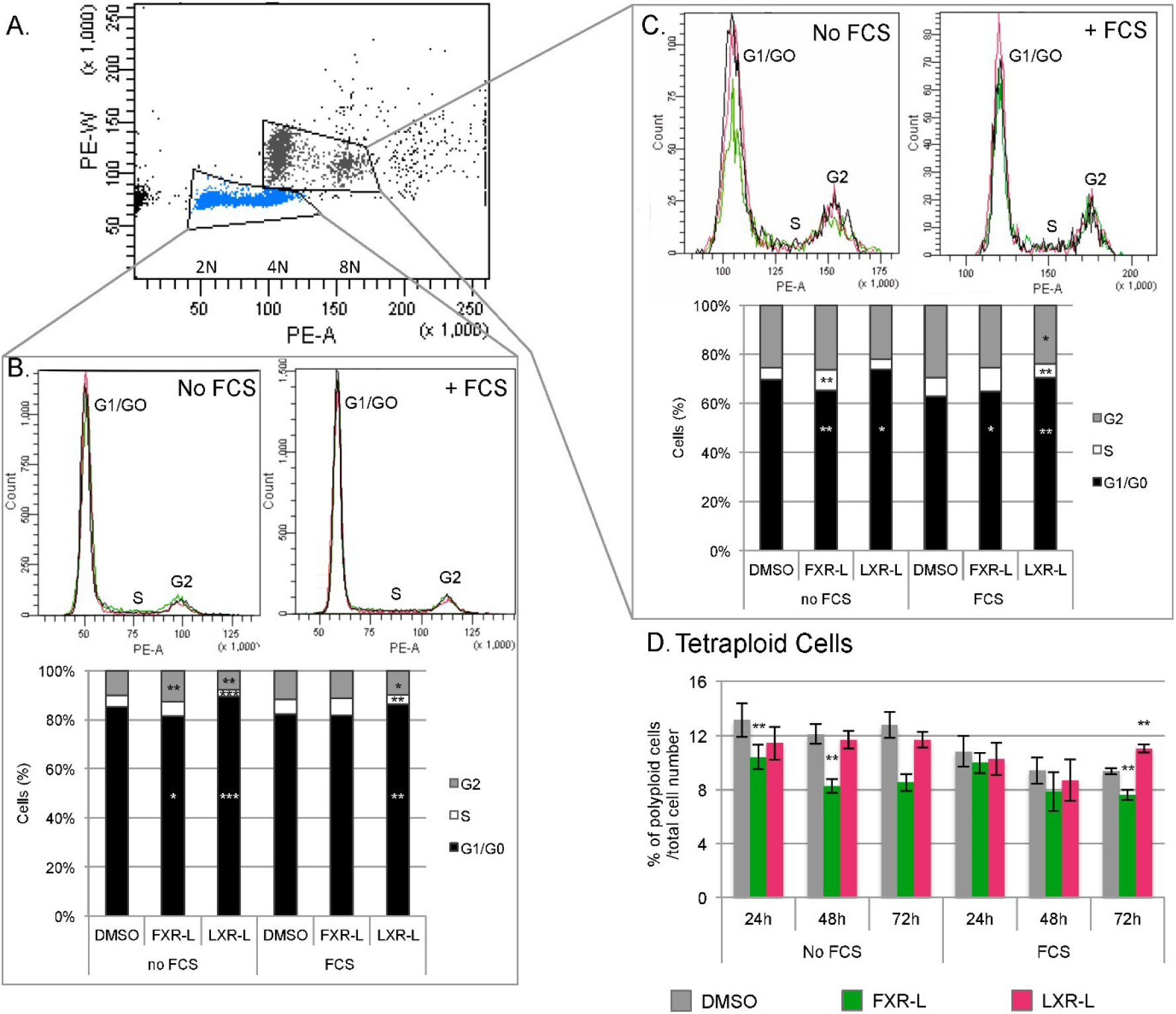
FXR and LXR affect the cell progression and the ploidy of HepaRG cells. The cell cycle distribution and the ploidy were analyzed by PI staining and FACS. A. The panel shows the group of HepaRG cells, the diploid cells (2N) and tetraploid cells (4N), B and C. The panels and the graphs show the cell cycle distribution at 72h of treatment with DMSO, FXR-L 1 uM or LXR-L 2 uM. D. The graph shows the percentage of tetraploid cells at 24h, 48h and 72h with the different treatments. *: p-value < 0.05, ** < 0.01, *** < 0.001 versus DMSO (Student’s t test). N=8.

The mechanisms involved in the mammalian hepatocyte polyploidism are not yet well understood. However, polyploidy is known to appear at the terminal differentiation during late foetal development, while proliferating cells are mainly diploid. By sorting the cells by their cell cycle phase status, we observe in the diploid population an accumulation of cells at G1 after 48 hours treatment with LXR-L (suppl. Fig. S5). This accumulation is further increased at 72 hours, while S and G2 populations are decreasing (Fig. 4B). In opposite manner, FXR-L tends to decrease the G1 population and increase the S and G2 populations at 24h, even if these changes are statistically significant only at 72h in serum-free conditions (Fig. 4B and suppl. Fig. S5). The tetraploid population shows the same pattern with a stronger effect of FXR-L, decreasing significantly the G1 population in both conditions (Fig. 4C). Hence, our results are consistent with a correlation between polyploidy and hepatocyte differentiation stage (31): FXR-L has a pro-proliferative effect and decreases polyploidy, while LXR-L inhibits the cell cycle progression, inducing hepatocyte differentiation and a higher polyploidism.

Positionning the genes identified in the GSEA results with the BIOCARTA pathways show that FXR-L affects the G1/S and G2/M checkpoints, while LXR-L only affects the G1/S checkpoint, blocking the cells in the G1 phase (Fig. 4). To better assess the key players involved in these regulations, we evaluated the activity of some cell cycle regulators at the transcriptional and post-transcriptional levels (Fig. 5). The LXR-L treatment affects the G1/S checkpoint by inhibiting the expression of *CDC25A* and *CCNE2* and up-regulating the expression *CDKN2B* (p15). In parallel, quantitative PCR shows a down-regulation of *E2F1* and *E2F2* mRNA expression by qPCR (Fig. 5A). The treatment also has effects at the post-transcriptional level inhibiting the CCND3 (Cyclin D3) expression and decreasing RB phosphorylation (Fig. 5B&C). FXR-L also deeply affects the key players of the G1/S checkpoint, *CCNE1* and *SKP2*, which are up-regulated by FXR-L (Fig. 5A). This is consistent with the observed hyper-phosphorylation of RB (Fig. 5B&C) and should result in an activation of the cell cycle progression. Finally, FXR-L also affects the mRNA expression of the G2/M checkpoint regulators *CDC25C* or *CCNA2* (Fig. 5A).

**Figure 5:**
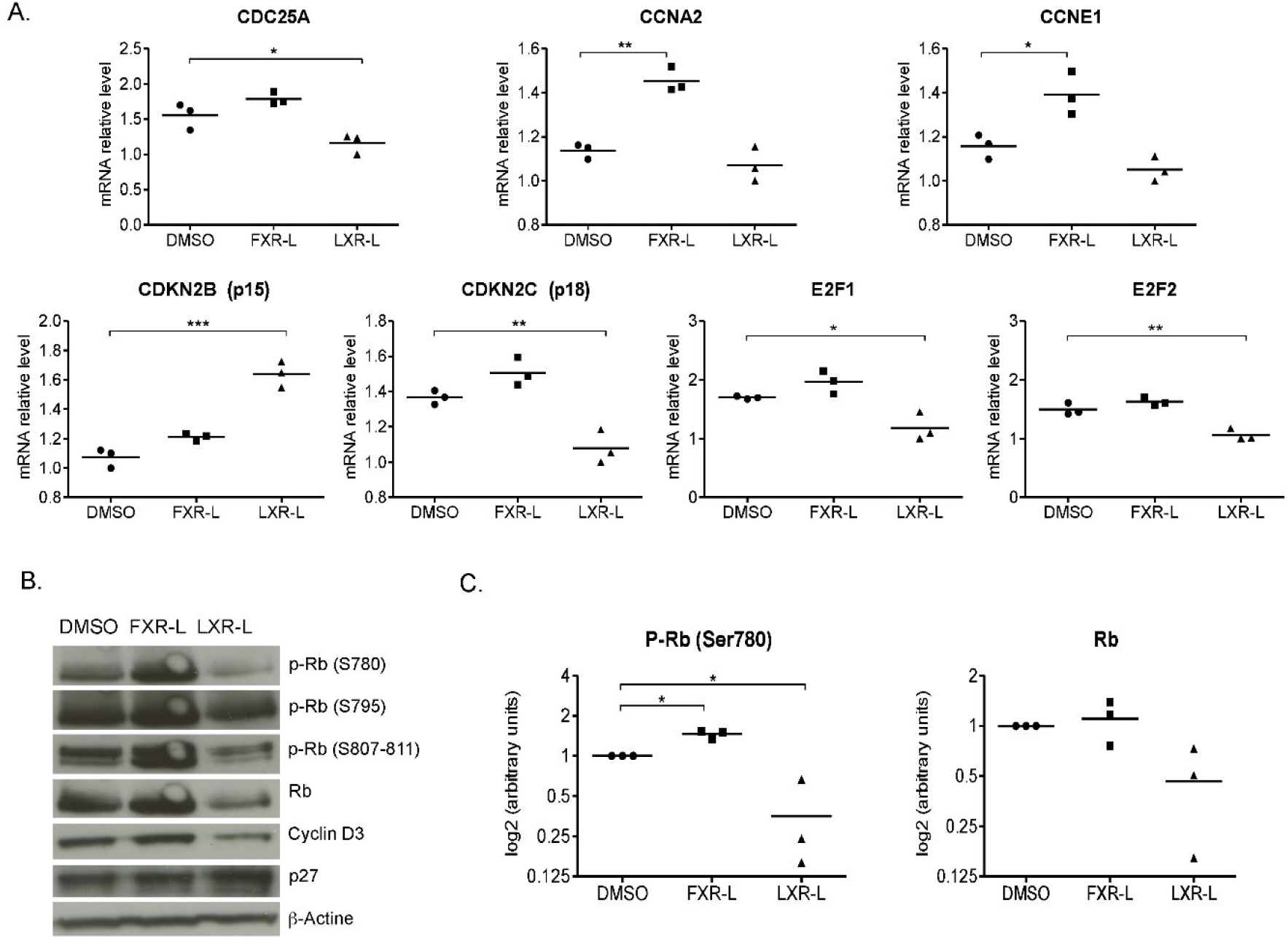
Validation of the effects of FXR and LXR on cell cycle regulators by RT-qPCR (A) and Western blot (B). HepaRG cells have been treated 24h in the same condition as for the microarrays (same ligands and DMSO as control). A. Relative mRNA levels for genes regulated in the microarray data. B. Representative Western blot of Retinoblastoma (Rb), phospho-Rb, Cyclin D3, and p27 in whole HepaRG cell lysates. C. densitometric quantification of phospho-Rb and Rb. *: p-value < 0.05, ** < 0.01 versus DMSO (Student’s t test). N=3.

## Discussion

Nuclear receptors are important transcriptional regulators of metabolism (2), acting on both specific and shared pathways. The numerous side effects associated to novel therapeutics for metabolic disorders based on NR activation or inhibition highlight the need for understanding their regulatory cross-talks. Herein, we demonstrated that the cross-talk between the three selected receptors, FXR, LXR and PPARα is largely due to a distribution of their respective specific target genes in shared pathways, rather than to their effects on common genes. This study further allowed identifying an important role of FXR and LXR in controlling, in opposite manner, some key factors of cell cycle progression in liver cells and of hepatocyte polyploidization.

One strength of this study is the use of HepaRG cells, which exhibit a gene expression profile close to the primary human cells. By using a simplified, well standardized, and human-related model, the results obtained are likely to be informative with respect to human responses to the ligands used herein. Another strength of this study was to obtain the profile of gene expression upon activation of FXR, LXR and PPARα, respectively, using the very same experimental conditions. This was the condition to be able to interpret the specificity of each receptor. Indeed, existing gene expression profiles for these receptors that have already been published in the GEO repository cannot be directly compared due to the different experimental setups and cell lines used for the studies (for example, HepG2 vs HepaRG, vs primary hepatocytes, see (5) and Suppl. Fig. S1).

Ideally, the cell culture system would have allowed to also test how NRs were acting on each other by knocking down one by one FXR, LXR and PPARα and evaluate how the deletion of one receptor would affect the specific response of another receptor. For that purpose, we had performed all the preliminary approaches (identification of the siRNA targeting the NRs of interest, construction of the adenovirus expressing these siRNAs, and validation of their capacity to inhibit their specific targets). However, the analyses of the first series of microarrays, following HepaRG infection with the adenovirus containing the control siRNA (targeting the non human gene Luciferase) or the specific siRNA targeting FXR, revealed that several pathways, in particular the one related to bile acid metabolism, were severely affected by the control adenovirus, precluding any rigourous analyses of the results with the specific siRNA. We consider that this unexpected outcome of adenovirus infection of liver cells has to be mentioned, for the sake of further research using this tool. More specifically, the possible alteration of the bile acid metabolism pathways should be checked when using adenovirus-related tools in liver cells or tissue.

An important observation in the present study is that the three receptors we tested regulate relatively few genes in common. There are several levels of promiscuousness between NRs: they share the same binding partner RXR, they have promiscuous binding sites, and they act on common metabolic pathways. However, our results emphasize their specificity in activating genes. This specificity does apply at the very early time point (4h), where we expect only direct target genes to be activated, but extend also to the late time point (24h), where new genes are activated, in part as a consequence of the first set of gene activation. Similar observations have been reported when analyzing the gene regulatory profile of PPARγ and LXR receptors in cancer cells. While they both exert an inhibitory action on cancer cell proliferation, via a common metabolic reprogramming, the genes they activate remain specific to each receptor (32).

Another feature of interest is the overlap but also specificity of the action of CDCA and FXR-L. CDCA is considered to be a natural ligand for FXR (11). However, a large number of genes are activated with the synthetic ligand but not with the natural one. This may be explained by the so-called selective NR modulator effect (33), by which the conformational changes that occur in NRs upon ligand binding may provoke a different interface for co-factor recruitment, depending on the nature of the ligand. It may also be caused by a different affinity of the ligand, a different accessibility, and/or different metabolism of the ligand within the cell. These facts are particularly important when searching for appropriate ligands aimed at therapeutic usage, implying that a full exploration of the gene expression profile for each new molecule proposed is required.

Unsurprisingly, the main pathways enriched by the three NR agonists are related to metabolism. More interestingly, 72% of the pathways shared by FXR and LXR are involved in regulations of cell growth and death. In addition, 56% of genes belonging to the leading edge of the cell cycle pathway (evaluated by GSEA) are common to both NRs but with an opposite modulation. The anti-proliferative activities of LXR has been observed in a number of cellular systems, including vascular smooth muscle cells, cancer cells, T lymphocytes, pancreatic islet beta cells and mouse liver (reviewed in (34)). This brought up the claim that LXR might be a target for anti-tumoral treatments in different tissues, mainly documented in breast, prostate and intestinal cancer cells. LXR mediated cell-cycle inhibition has been mechanistically correlated with the expression of lipogenic and triglyceride accumulation (28), but could also be related to the inhibition of SKP2 transcription (35). In the present study, SKP2 is not affected by LXR-L, suggesting that LXR’s anti-proliferative activity is due to alternative mechanisms, such as LXR-mediated repression of AP1 signaling (32, 36). The role of FXR is more controversial, or complex. FXR was demonstrated to be involved in the bile-acid dependent progression of intestinal metaplasia (37), possibly through a SHP-dependent increase of CDX2 gene expression (38). In contrast, the deletion or downregulation of FXR favors hepatocarcinogenesis (27, 39), suggesting that FXR plays a protective role. Finally, FXR is also required for liver regeneration (24, 40). The mechanism by which FXR can both favor liver regeneration and protect from hepatocarcinogenesis remains unsettled (39). The following three main findings of the present study are noteworthy in this context. First, FXR-L significantly upregulates IL6 and increases the expression of a number of pro-inflammatory cytokines, such as *IL1B*, *CSF1*, *CSF2*, and *TGFB3*. These signals are indeed important in triggering the early response for liver regeneration (39), even though these results are in contradiction with the reported anti-inflammatory activities of FXR, via down-regulation of the NF-KB pathway (41), not observed in our dataset. Second, we showed the positive effect of FXR on cell cycle progression, increasing the activities of key regulators of the G1/S and G2/M checkpoints. Third, we show that FXR-L inhibits hepatocyte polyploidization. This effect could involve the insulin-signaling pathway, which is sitting at the cross-road of many metabolic pathways, and contributes to the formation of binucleated tetraploid liver cells through the PI3K/Akt pathway (42). Indeed, the insulin pathway is affected by all treatments in this study, but more particularly by FXR-L at 24h (p-value<0.001), with an inhibition of 17 key genes including INSR, IRS and PI3K (Suppl. Fig. S4). This suggests that FXR may prevent the polyploidization of the HepaRG cells via inhibition of the insulin INSR/PI3K pathway. Interestingly, polyploidy was shown to trigger cell transformation and tumor formation in p53-null cells (43). Although it remains difficult to generalize this observation to other cellular context, the inhibition of polyploidization mediated by FXR-L may contribute to its protective role against carcinogenesis.

Thus, besides illustrating the high levels of target gene specificity of each NR tested herein, our results revealed an important antagonistic effect of FXR and LXR on cell cycle progression in hepatocytes. Assessing these roles in different cells and tissues might be of importance when contemplating the therapeutic applications that these two receptors are conveying.

## Acknowledgements

We thank Frédéric Schütz and Thierry Sengstag for making their unpublished R package for gene set enrichment analysis (mygsea2) available, and we are grateful to Frédéric Schütz for excellent input on how to choose parameter settings. Particular thanks go to Christiane Guguen-Guillouzo and her laboratory for the generous gift of HepaRG cells and shared information on their maintainance. The Genomic Technologies Facility of the University of Lausanne has realized all the microarrays including data normalization.

