## Supplemental figures for "System analysis of cross-talk between nuclear receptors reveals an opposite regulation of the cell cycle by LXR and FXR in human HepaRG liver cells"

**In addition to these supplementary figures, a file will be provided only as numerical material. This file is referred to in the text as an archive "data_and_code.zip". It will contains the detailed list of genes, pathways, and categories.**

**Figure legends of supplementary figures**

**Figure S1: Hierarchical clustering of 15 microarray samples of liver cells and various hepatic cell lines (publicly available Affymetrix data set from Hart et al. (reference to bibliography).** Sample clustering was based on the expression profiles of the 1750 genes that were found significantly regulated by any of the four NR-treatments (FXR-L, CDCA, LXR-L, PPARA-L) in our own microarray study at either of the two time points (4h, 24h). As expected, primary hepatocytes (phh) are the closest to liver cells. Next closest are the HepaRG cells, while the HepG2 cells are clustering apart from all other cell types.
Significance testing for gene selection: Moderated F-tests with four NR-vs-control contrasts were performed separately per time point; p-values were adjusted by the Benjamini-Hochberg method; and genes with adjusted $p < 0.05$ in one or both of the two result lists were retained for the clustering analysis. Clustering method: Ward's minimum variance method, as implemented in the R function hclust prior to R version 3.0.3.

**Figure S2: Illustration of the GSEA method and selection of leading edge genes,** see methods for more details. The horizontal axis represents genes, sorted according to their p-value in LIMMA (shown in the bottom panel). Genes associated to the pathway of interest (here the cell cycle) correspond to the vertical stripes in the center. On the top part, the blue curve represents the GSEA score, which increases when the current gene is part of the pathway and decreases otherwise (increasing and decreasing steps depend on the number of genes remaining to visit). The red vertical line marks the maximal score reached. The leading edge genes are the ones which have been visited before this line. The genes in over-represented (resp. under-represented) pathways have a higher density in the start (resp. end) of the list, which drives a high positive (resp. negative) score. The significance of these scores is assessed by permutations (see methods).

**Figure S3: Effect of the FXR-L and CDCA treatments on the KEGG cell cycle pathway at 4h.**

**Figure S4: Effect of the FXR-L and LXR-L treatments on the KEGG Insulin pathways. A**) The genes in the leading edge for FXR-L treatment at 24h are hightlighted, as in main figure 3 (blue for down-regulations, orange for up-regulations). **B)** Same image for the LXR-L treatment at 24h.

**Figure S5: FXR and LXR affect the cell progression and the ploidy of HepaRG cells.**
The cell cycle distribution and the ploidy were analyzed by PI staining and FACS. A and B show the cell cycle distribution at 24h of treatment with DMSO, FXR-L 1 uM or LXR-L 2 uM. C and D show the cell cycle distribution at 48h with the different treatments. * : p-value < 0.05, ** < 0.01, *** < 0.001 versus DMSO (Student's t test). N=8.

Figure S1

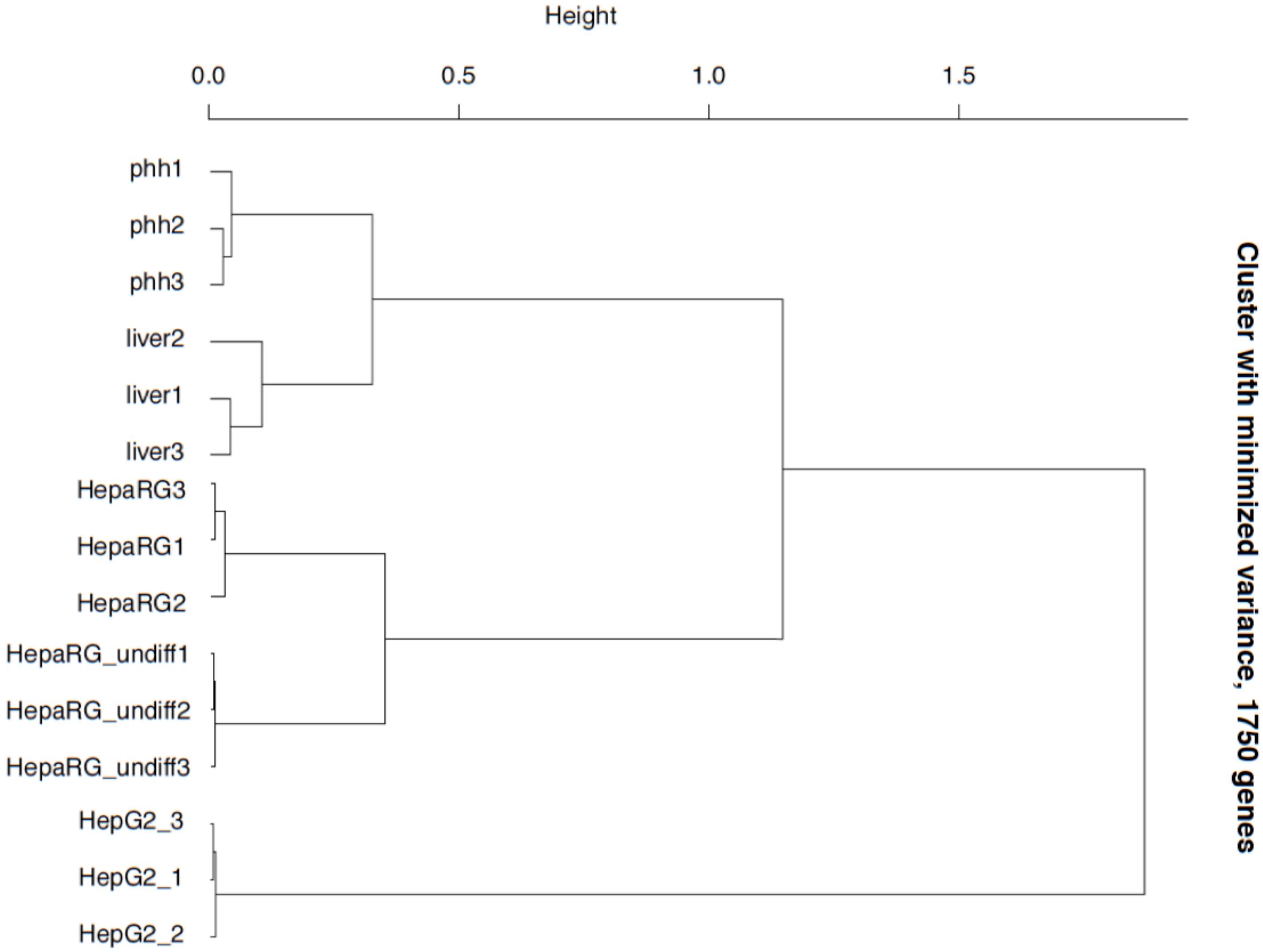

Height
0.0
0.5
1.0
1.5
phh1
phh2
phh3
liver2
liver1
liver3
HepaRG3
HepaRG1
HepaRG2
HepaRG_undiff1
HepaRG_undiff2
HepaRG_undiff3
HepG2_3
HepG2_1
HepG2_2
Cluster with minimized variance, 1750 genes

Figure S2

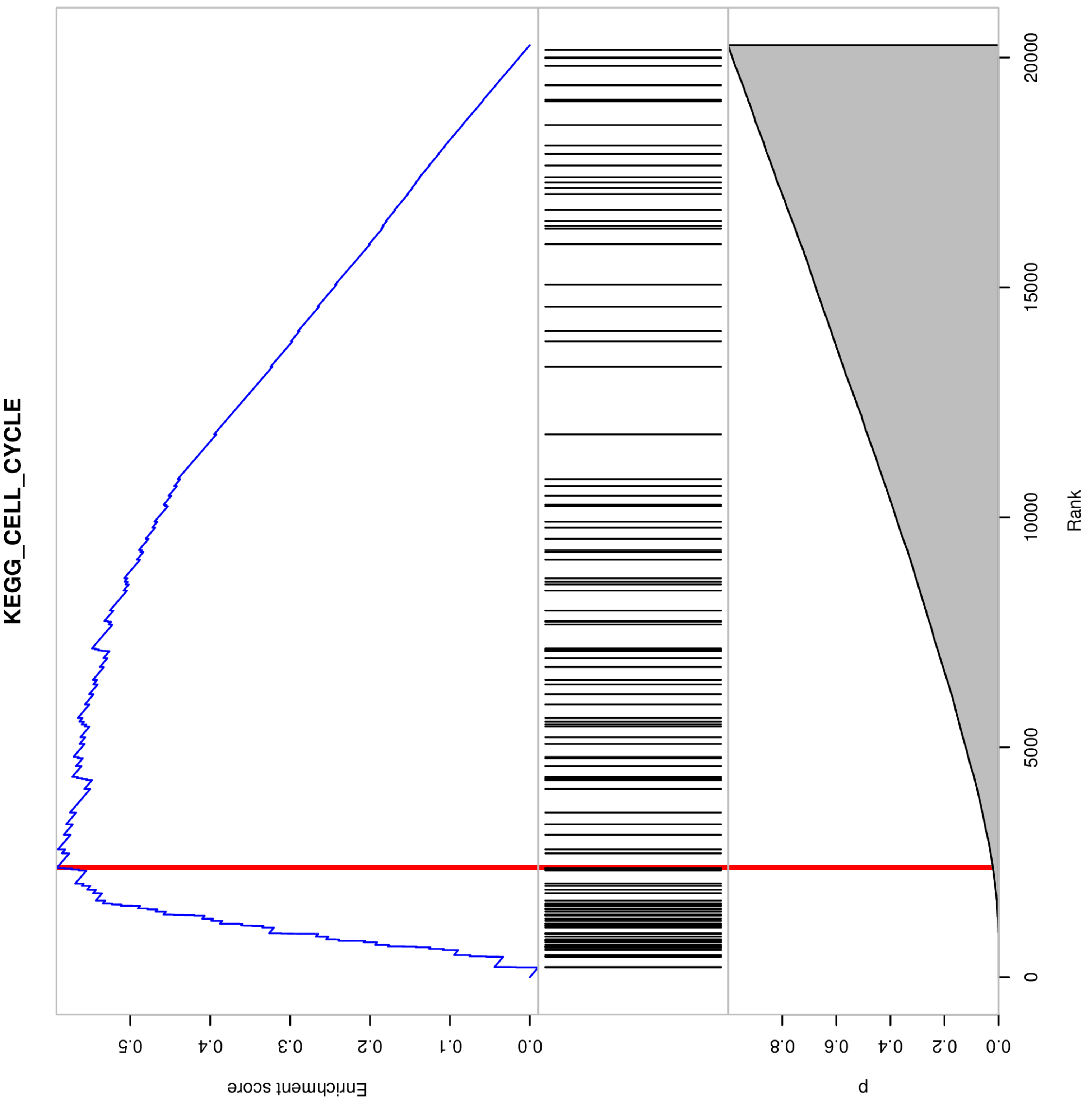

KEGG_CELL_CYCLE
Enrichment score
0.0
0.1
0.2
0.3
0.4
0.5
p
0.0
0.2
0.4
0.6
0.8
0
5000
10000
15000
20000
Rank

Figure S3a

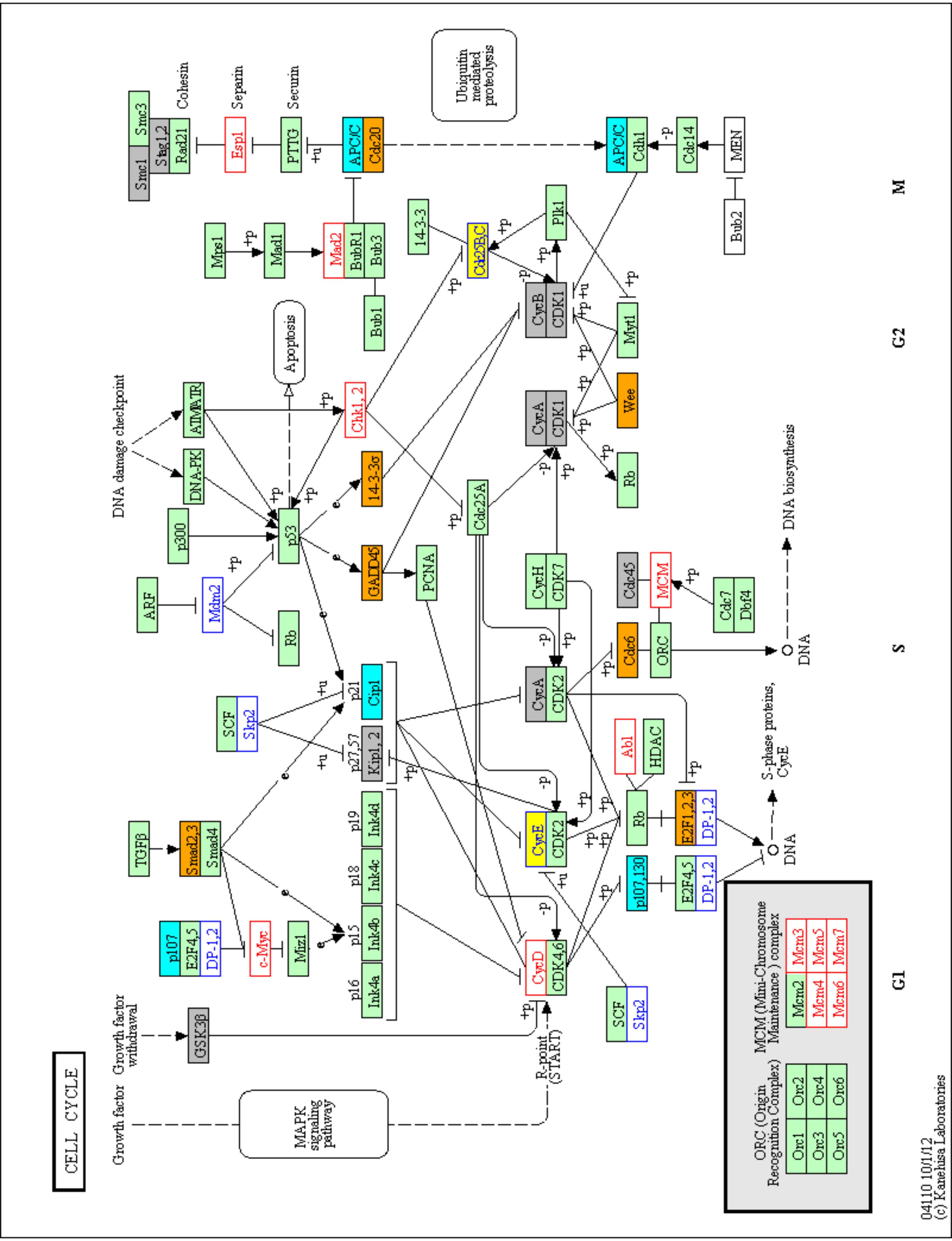

CELL CYCLE
Growth factor
Growth factor withdrawal
MAPK signaling pathway
R-point (START)
GSK3β
TGFβ
Smad2,3
Smad4
p107
E2F4,5
DP-1,2
c-Myc
Miz1
p16
Ink4a
p15
Ink4b
p18
Ink4c
p19
Ink4d
p27,57
Kip1, 2
p21
Cip1
SCF
Skp2
ARF
Mdm2
Rb
p300
p53
DNA-PK
ATM/ATR
DNA damage checkpoint
Apoptosis
Chk1, 2
14-3-3σ
GADD45
PCNA
Mps1
Mad1
Mad2
BubR1
Bub3
Bub1
Smc1
Smc3
Stag1,2
Rad21
Cohesin
Esp1
Separin
PTTG
Securin
APC/C
Cdc20
Cdh1
Ubiquitin mediated proteolysis
14-3-3
Cdc25B,C
Plk1
CycB
CDK1
CycA
CDK1
Cdc25A
CycH
CDK7
CycA
CDK2
CycE
CDK2
CycD
CDK4,6
Myt1
Wee
Cdc45
MCM
Cdc6
ORC
Cdc7
Dbf4
Abl
HDAC
E2F1,2,3
p107,130
Cdc14
MEN
Bub2
DNA biosynthesis
S-phase proteins, CycE
DNA
ORC (Origin Recognition Complex)
Orc1
Orc2
Orc3
Orc4
Orc5
Orc6
MCM (Mini-Chromosome Maintenance ) complex
Mcm2
Mcm3
Mcm4
Mcm5
Mcm6
Mcm7
G1
S
G2
M
04110 10/1/12
(c) Kanehisa Laboratories

Figure S3b

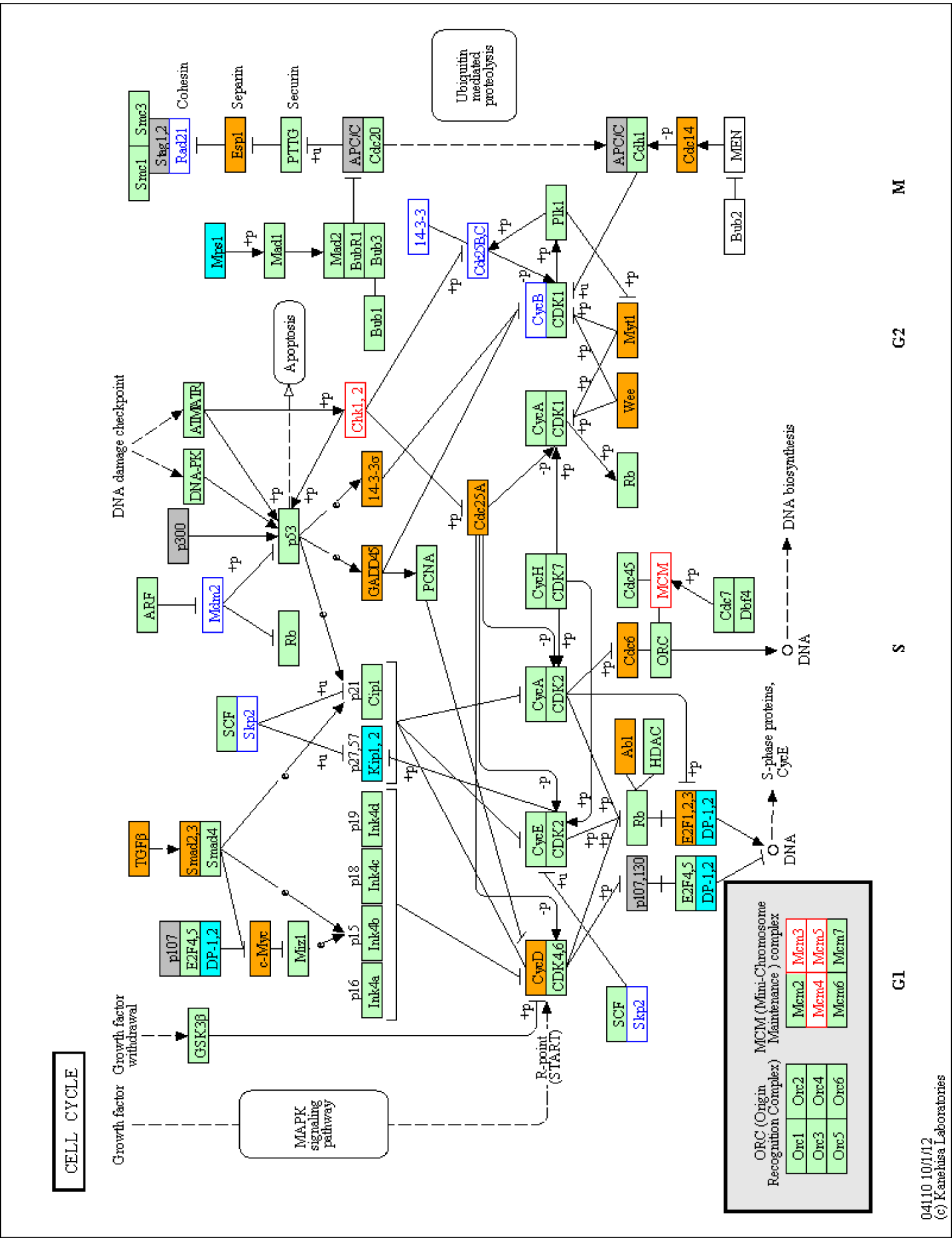

CELL CYCLE
Growth factor
Growth factor withdrawal
MAPK signaling pathway
GSK3β
R-point (START)
DNA damage checkpoint
ATM/ATR
DNA-PK
p300
p53
Chk1, 2
14-3-3σ
GADD45
PCNA
Apoptosis
ARF
Mdm2
Rb
TGFβ
Smad2,3
Smad4
p107
E2F4,5
DP-1,2
c-Myc
Miz1
SCF
Skp2
p21
Cip1
p27,57
Kip1, 2
p19
Ink4d
p18
Ink4c
p15
Ink4b
p16
Ink4a
CycD
CDK4,6
CycE
CDK2
CycA
CDK2
CycH
CDK7
Cdc25A
CycA
CDK1
CycB
CDK1
Cdc25B,C
14-3-3
Plk1
Wee
Myt1
Mps1
Mad1
Mad2
BubR1
Bub3
Bub1
Smc1
Smc3
Stag1,2
Rad21
Cohesin
Esp1
Separin
PTTG
Securin
APC/C
Cdc20
Cdh1
Cdc14
MEN
Bub2
Ubiquitin mediated proteolysis
Abl
HDAC
E2F1,2,3
p107,130
Cdc45
MCM
Cdc6
ORC
Cdc7
Dbf4
DNA
DNA biosynthesis
S-phase proteins, CycE
ORC (Origin Recognition Complex)
Orc1
Orc2
Orc3
Orc4
Orc5
Orc6
MCM (Mini-Chromosome Maintenance ) complex
Mcm2
Mcm3
Mcm4
Mcm5
Mcm6
Mcm7
G1
S
G2
M
04110 10/1/12
(c) Kanehisa Laboratories

Figure S4

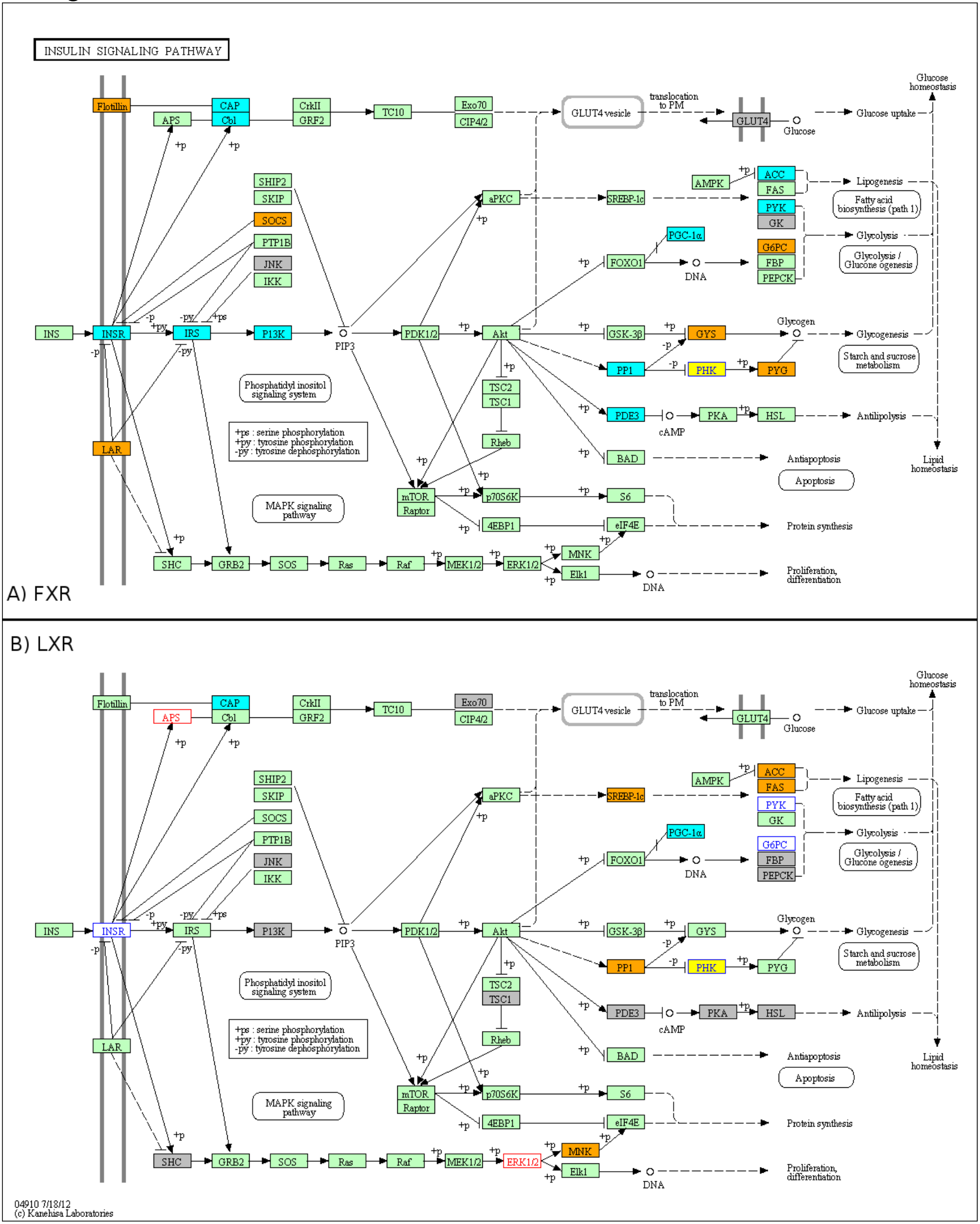

INSULIN SIGNALING PATHWAY
A) FXR
B) LXR
INS
INSR
IRS
PI3K
PIP3
PDK1/2
Akt
aPKC
Flotillin
APS
CAP
Cbl
CrkII
GRF2
TC10
Exo70
CIP4/2
GLUT4 vesicle
translocation to PM
GLUT4
Glucose
Glucose uptake
Glucose homeostasis
SHIP2
SKIP
SOCS
PTP1B
JNK
IKK
LAR
SHC
GRB2
SOS
Ras
Raf
MEK1/2
ERK1/2
MNK
Elk1
DNA
Proliferation, differentiation
SREBP-1c
AMPK
ACC
FAS
Lipogenesis
Fatty acid biosynthesis (path 1)
PYK
GK
Glycolysis
PGC-1α
FOXO1
G6PC
FBP
PEPCK
Glycolysis / Gluconeogenesis
GSK-3β
GYS
Glycogen
Glycogenesis
PP1
PHK
PYG
Starch and sucrose metabolism
TSC2
TSC1
Rheb
PDE3
cAMP
PKA
HSL
Antilipolysis
Lipid homeostasis
BAD
Antiapoptosis
Apoptosis
mTOR
Raptor
p70S6K
S6
4EBP1
eIF4E
Protein synthesis
Phosphatidyl inositol signaling system
MAPK signaling pathway
+ps : serine phosphorylation
+py : tyrosine phosphorylation
-py : tyrosine dephosphorylation
04910 7/18/12
(c) Kanehisa Laboratories

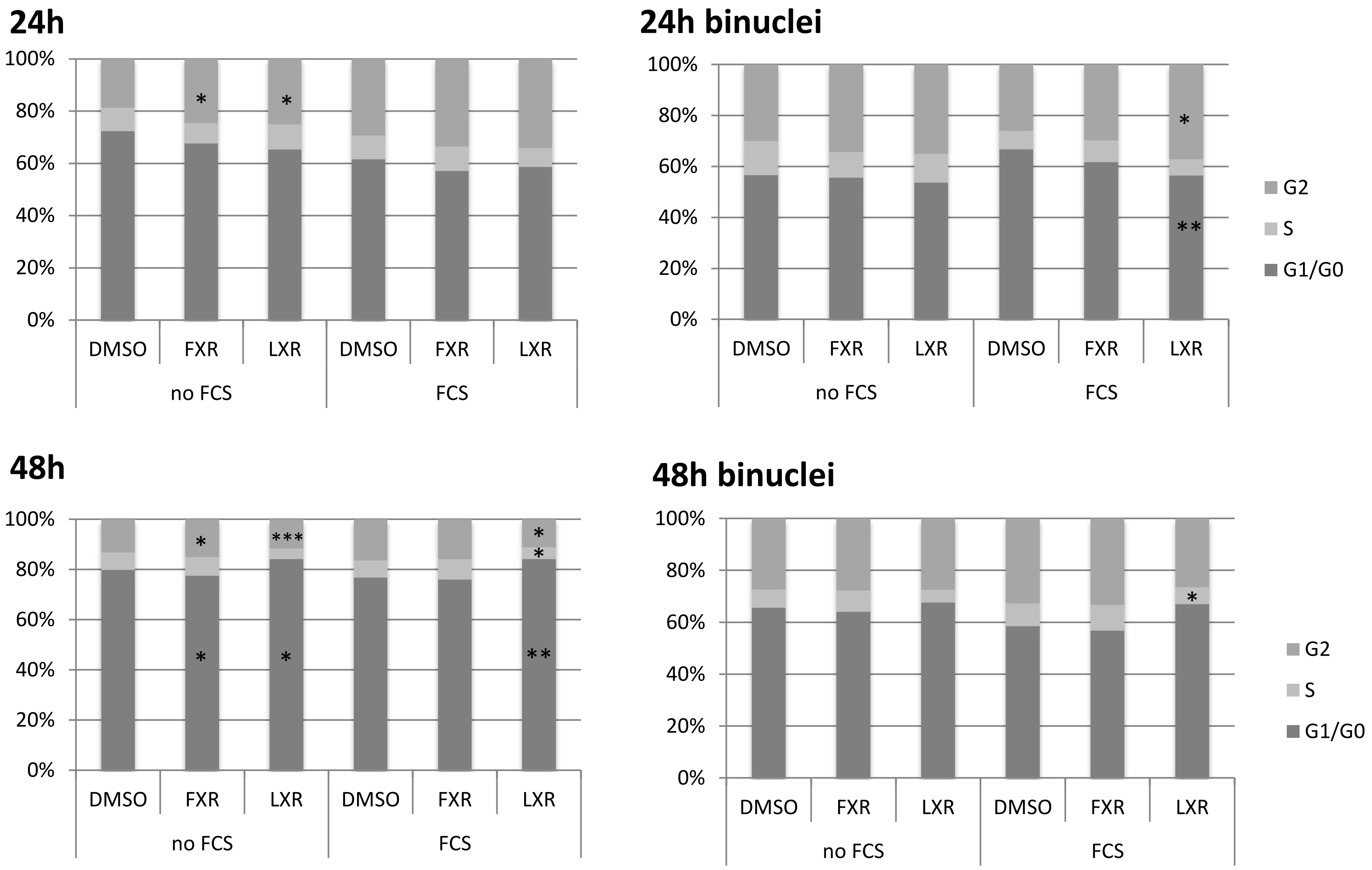
24h
24h binuclei
48h
48h binuclei
100%
80%
60%
40%
20%
0%
DMSO
FXR
LXR
no FCS
FCS
G2
S
G1/G0
*
**
***

Figure S5
