## Supplemental data and code for "System analysis of cross-talk between nuclear receptors reveals an opposite regulation of the cell cycle by LXR and FXR in human HepaRG liver cells": count_genes_pathways.ipynb

```
# %% [code]
import os
import os.path
import itertools
import pandas as pd

ligands = ('CDCA', 'FXR', 'LXR', 'PPARA')
fxr_ligands = ('FXR+', 'LXR', 'PPARA')
timepoints = (4,24)

# %% [markdown]
# Helper methods

The following methods parse the input files and extract sublists.

# %% [code]
def load_genes(ligands, timepoints, t_enrich):
    raw_data = {}
    for l in ligands:
        for tp in timepoints:
            filename = os.path.join('genes','%s_%sh.txt' % (l,tp))
            sub = pd.read_table(filename)[["ID", "adj.P.Val", "logFC"]]
            sub = sub[sub["adj.P.Val"]<t_enrich]
            print("LOAD: [%s,%s] -> %s" % (l,tp, len(sub)))
            raw_data[(l,tp)] =  { row.ID: row.logFC  for idx,row in sub.iterrows() }
    return raw_data

def load_kegg_pathways(ligands, timepoints, t_enrich):
    raw_data = {}
    for l in ligands:
        for tp in timepoints:
            filename = os.path.join('pathways', 'mygsea2_%s_%sh.txt' % (l,tp))
            t = pd.read_table(filename)
            sub = t[["geneset", "p.pos.adj"]]
            sub = sub[sub["p.pos.adj"]<t_enrich]
            ksub = sub[ sub["geneset"].str.startswith("KEGG")]
            raw_data[(l,tp)] = set( [ n for n in ksub["geneset"]] )
    return raw_data

def cross(ligand_lists, categories=None):
    for l in ligand_lists:
        print('%s:' % l)
        data = ligand_lists[l]
        total = len(data)
        single = data
        others = [d for d in ligand_lists.keys() if d != l]
        for o in others:
            d = ligand_lists[o]
            single = single.difference(d)
        show_group('single', single, categories)

        multiple = data.difference(single)
        for o in others:
            crossed = data.intersection(ligand_lists[o])
            for oo in others:
                if oo != o:
                    crossed = crossed.difference( ligand_lists[oo] )
            show_group( "cross %s" % o, crossed, categories)
            multiple = multiple.difference(crossed)
        show_group('Multiple', multiple, categories)
        show_group('Total', data, categories)

def show_group(name, data, categories):
    print("    %s: %s" % (name, len(data)), end='')
    if categories:
        print(' -> ', end='')
        for c in categories:
            catdata = data.intersection(categories[c])
            print('%s ' % len(catdata), end=' ')
    print()

def table1(raw_data, ligands=ligands, categories=None):
    ligand_lists = {}
    time_lists = {}
    for t in (4,24):
        time_lists[t] = set()
    for l in ligands:
        llist = set()
        for t in timepoints:
            cur = raw_data[(l,t)]
            time_lists[t] = time_lists[t].union(cur)
            llist = llist.union(cur)
        ligand_lists[l] = llist

    cross(time_lists, categories)
    print()
    cross(ligand_lists, categories)

def load_categories():
    categories = {}
    for name in ('metabolism', 'cell_growth_death', 'immune') :
        f = open( os.path.join('categories', '%s.txt' % name))
        members = set()
        for l in f.readlines():
            l = l.strip()
            members.add(l)
        categories[name] = members
    return categories

# %% [markdown]
# Distribution of genes and pathways (Table 1)

# %% [code]
raw_genes = load_genes(ligands, timepoints, 0.05)
table1( raw_genes )

# %% [code]
raw_pathways = load_kegg_pathways(ligands,timepoints, 0.2)
table1(raw_pathways)

# %% [markdown]
# Categories of pathways (Table 2)

# %% [code]
# Keeping CDCA and FXR separate
categories = load_categories()
table1(raw_pathways, categories=categories)

# %% [code]
# Merging CDCA and FXR
fxr_pathways = raw_pathways.copy()
for t in timepoints:
    del( fxr_pathways[('FXR',t)] )
    del( fxr_pathways[('CDCA',t)] )
    fxr_pathways[("FXR+",t)] = raw_pathways[('FXR',t)].union(raw_pathways[('CDCA',t)])

table1(fxr_pathways, ligands=fxr_ligands, categories=categories)

# %% [markdown]
# Construct all groups for loading into cytoscape (Fig 1)

# %% [code]
conditions = [
    ('c', ('CDCA', 4)),
    ('f', ('FXR',  4)),
    ('l', ('LXR',  4)),
    ('p', ('PPARA',4)),
    ('C', ('CDCA', 24)),
    ('F', ('FXR',  24)),
    ('L', ('LXR',  24)),
    ('P', ('PPARA',24)),
]

"Assemble all lists of selected genes or pathways into a single dataframe"
def load_global(data, hasdata=False):
    rows = set()
    for c, ctp in conditions:
        rows.update(data[ctp])
    df = pd.DataFrame(index=rows)
    ddf = None
    if hasdata:
        ddf = pd.DataFrame(index=rows)
    for c,ctp in conditions:
        df[c] = pd.Series( index=data[ctp], data=True)
        if hasdata:
            ddf[c] = pd.Series( data=data[ctp])
    df.fillna(False, inplace=True)
    if hasdata:
        ddf.fillna(False, inplace=True)
    return df,ddf

def make_groups(df, ddf, outpattern, cnt_th=25):
    out_nodes = open('%s_nodes.tsv' % outpattern, 'w')
    out_nodes.write('uid\tname\ttype\tweight\n')
    out_edges = open('%s_edges.tsv' % outpattern, 'w')
    out_edges.write('uid\tsource\ttarget\tweight\tratio\n')

    for c,ctp in conditions:
        out_nodes.write('%s\t%s %s\t%s\t\n' % (c, ctp[0], ctp[1], ctp[0]))
    links = []

    titles = [ c for c in df.columns ]
    l = len(titles)

    grouped = df.groupby( titles )
    cur_ddf = None
    for g in grouped:
        c = g[0]
        cnt = len(g[1])
        uid = "".join([ titles[i] for i in range(l) if c[i] ])
        if cnt >= cnt_th:
            name = cnt
        else:
            name = ""
        out_nodes.write('g_%s\t%s\tgroup\t%s\n' % (uid, name, cnt))
        
        
        idxs = tuple( g[1].index )
        ratio = .5
        if ddf is not None:
            cur_ddf = gfc_df.loc[ idxs, ]
        for cnd in uid:
            if cur_ddf is not None:
                # Compute the positive/negative ratio
                tmp = gfc_df.loc[ idxs, (cnd)]
                ratio = len(tmp[ tmp>0]) / len(tmp)
            out_edges.write("%s_%s\t%s\tg_%s\t%s\t%s\n" % (cnd,uid,cnd,uid,cnt, ratio))

    out_nodes.close()
    out_edges.close()

# %% [code]
g_df,gfc_df = load_global(raw_genes, True)
make_groups(g_df, gfc_df, os.path.join('output', 'grp_genes'))

p_df,pfc_df = load_global(raw_pathways)
make_groups(p_df, pfc_df, os.path.join('output', 'grp_pathways'), cnt_th=4)
```
